# A Computational Framework for Analysis of cfDNA Fragmentation Profiles

**DOI:** 10.1101/2025.04.17.649313

**Authors:** Zhong Wee Poh, Guanhua Zhu, Pui Mun Wong, Denis Odinokov, Hanae Carrie, Yi Ting Lau, Anna Gan, Polly Poon, Patrick Tan, Anders J. Skanderup, Iain Tan

## Abstract

Circulating cell-free DNA (cfDNA) has emerged as a promising non-invasive medium for studying tumor molecular profiles. Non-random fragmentation patterns in plasma cfDNA, particularly around nucleosome-depleted regions (NDRs) near transcription start sites (TSS), have been shown to reflect epigenetic regulation and gene expression. In this study, coverage profiles of the NDR were utilized to derive an NDR score, which was subsequently used as a proxy for inferring gene expression. To reduce transcript-to-transcript variability and enhance the clarity of these expression-associated signals, we implement a method for GC-bias correction of cfDNA samples. A computational framework (NDRDiff) was then developed to enable comparative analyses of NDR score profiles across different sample groups.

The GC-bias correction preserved the overall trend of the NDR signal while improving the separation of gene expression levels, as demonstrated by comparisons of healthy donor cfDNA samples with matched blood RNA-seq data. Validation on a simulated dataset showed that NDRDiff achieved an area under the precision–recall curve (AUPRC) of 0.916, outperforming a standard t-test (AUPRC of 0.777).

When applied to a comparison of healthy donor cfDNA and metastatic colorectal cancer (mCRC) cfDNA, NDRDiff identified 531 differential NDR score (DNS) genes that facilitated clear separation between the two groups. These DNS genes were found to correlate with tumor fraction estimates (down-regulated DNS genes: Pearson R = 0.89, p < 0.05; up-regulated DNS genes: Pearson R = –0.88, p < 0.05) and included CLDN4, BIN2, and IRAG2, which exhibit strong associations with colorectal cancer or blood cell expression signatures. Gene set enrichment analysis further revealed enrichment of colon and other gastrointestinal tissue signatures. Collectively, these findings underscore the potential of NDR-based cfDNA analysis as a minimally invasive tool for monitoring tumor-related molecular features in cancer.

## Introduction

Colorectal cancer (CRC) is one of the most prevalent cancers globally, accounting for about 10% of all diagnosed cancers and is the second leading cause of cancer-related deaths^1^. Approximately 22% of newly diagnosed CRC patients present with distant metastases^2^, and up to 70% of CRC patients develops metastatic disease^3^. Although advances in treatment have improved outcomes for selected patients, survival rates for metastatic CRC (mCRC) remains limited, with a 5-year relative overall survival (OS) of approximately 15%^4,5^. This underscores the importance of comprehensive management strategies, including tailored treatments based on molecular characteristics, to improve clinical outcomes for patients.

Understanding the evolution of the tumor molecular profile throughout the treatment period is critical for understanding the mechanisms of treatment resistance, enabling the early detection of resistance, and identifying opportunities for alternative therapies. However, this necessitates longitudinal sampling at multiple time points during treatment. Traditional methods for detecting these alterations often require invasive tissue biopsies, which are not only poses significant risk for patients but also may not capture the full heterogeneity of the tumor^6^. A liquid biopsy approach, such as the study of plasma cell-free DNA (cfDNA), presents an alternative to tissue biopsy due to the minimally invasive nature, allowing longitudinal sampling to track tumor evolution. Plasma cfDNAs are DNA released from cells into the blood and are released through diverse cell-death modalities, such as apoptosis and necrosis and active excretion mechanisms^7^. While predominantly originating from hematopoietic cells^8^, elevated plasma cfDNA is detected in cancer patients in 1977^9^ and since then, numerous studies have confirmed the contribution of tumor-derived DNA to circulating cfDNA^10–12^, referred to as circulating tumor DNA (ctDNA). Plasma ctDNA can retain genetic and epigenetic profile of tumor and has been used to detect genetic mutations^9,13,14^ and epigenetic alterations, such as DNA methylation^11,12,15^ and histone modification^16^.

Plasma cfDNA was found to show non-random fragmentation pattern that reflects epigenetic regulation^17^. Subsequently, differences in nucleosome protection patterns relative to the transcription start site (TSS) are observed between highly and lowly expressed genes. This pattern, or nucleosome footprint, around the TSS reflects chromatin accessibility possibly arising from the presence of the transcription preinitiation complex in transcriptionally active genes^18^. Later studies confirmed the finding and showed that the relative coverage of the nucleosome-depleted region (NDR) positioned from −150 to +50 bp relative to the TSS (henceforth referred to as NDR score) can be employed for the classification of expressed and silent genes^19^, and for the estimation of plasma cfDNA tumor fraction^20^. Considering the NDR score’s association with gene expression, we investigated the potential of using NDR score as a metric to infer gene expression profiles of the cellular populations contributing to plasma cfDNA.

In this study, we adapted existing GC-bias estimation approach to correct GC-bias in cfDNA samples. Building on these enhanced NDR estimates, we develop a computational framework for comparative analysis of NDR score profiles across different sample groups. We evaluate the performance of this framework against standard statistical test by validating it on a simulated dataset, and further demonstrate its utility by comparing healthy donor cfDNA samples with those from mCRC patients.

## Results

Deep whole-genome sequencing (dWGS) of plasma cfDNA was performed, achieving a sequencing coverage of 60x for healthy donors and 120x for mCRC samples (Methods). In total, the study included 10 mCRC patients, with 31 mCRC dWGS samples, and 17 dWGS samples from healthy donors (Table 1).

**Table 1.** Clinical characteristics of the dWGS cohort, including healthy and mCRC samples.

|  | Healthy (n=17) | mCRC (n=31) |
| --- | --- | --- |
| **<u>Age</u>** |  |  |
| Median (Min - Max) | 39.5 (23 - 52) | 57.2 (40.2 - 68) |
| **<u>Gender</u>** |  |  |
| Male | 2 | 7 |
| Female | 10 | 3 |
| **<u>Microsatellite Stability</u>** |  |  |
| **<u>Clinical</u>** |  |  |
| MSS | NA | 7 |
| Not Available | NA | 3 |
| **<u>Stage</u>** |  |  |
| III | NA | 1 |
| IV | NA | 9 |
| **<u>Tumor Location</u>** |  |  |
| Left Colon | NA | 4 |
| Right Colon | NA | 2 |
| Rectum | NA | 4 |
| **<u>Tumor Fraction</u>** |  |  |
| Median (Min - Max) | NA | 0.225 (0 - 0.506) |

### Inference of gene expression with NDR Score

As hematopoietic cells are the main contributors of plasma cfDNA in healthy individuals^21^, we investigated the association between NDR score and gene expression by comparing the relative coverage of TSS in our healthy samples (n=17) with corresponding gene expression from RNA-Seq data of blood samples.

Transcripts were categorized based on transcripts per million (TPM) values extracted from GTEx Whole Blood RNA-Seq. Comparing the mean relative coverage profiles across each TPM group, we demonstrated a reduction in relative coverage surrounding the TSS region associated with increased gene expression levels (Figure 1b), consistent with previous studies^19^. Specifically, we observed a decrease in mean relative coverage within the central region (−150 to +50 bp with respect to TSS) in actively transcribed genes compared to silenced genes.

**Figure 1.**
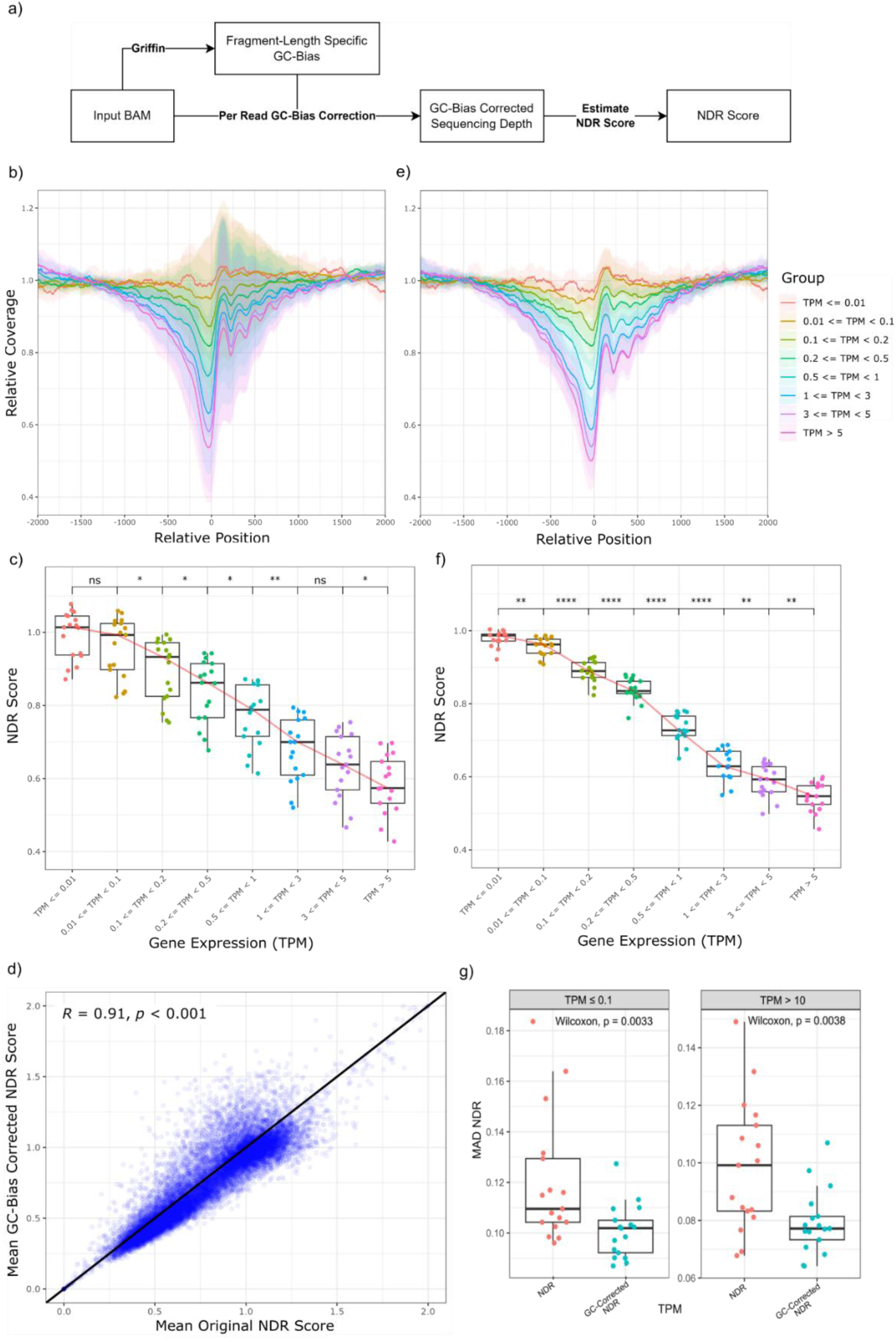
Association of NDR score with gene expression and improvement of NDR signals with GC-bias correction. **(a)** Schematic of GC-bias correction procedure for NDR score estimation. **(b, e)** Mean relative coverage around the TSS grouped for (b) original and (e) GC-bias corrected sequencing coverage according to gene expression (TPM) groups based on GTEx Whole Blood RNA-Seq for healthy cfDNA samples (n=17). The ribbon indicates the mean relative coverage ± 2 standard deviation (SD). The NDR site region is indicated by the dotted lines (−150 to +50 bp). **(c, f)** Comparison of mean NDR score across blood gene expression groups for (c) original and (f) GC-bias corrected NDR scores. **(d)** GC-bias corrected (y-axis) compared to original mean NDR scores (x-axis). **(g)** Comparison of mean absolute deviation (MAD) of original vs GC-bias corrected NDR score for silenced (TPM ≤ 0.1) and expressed genes (TPM > 10).

Subsequently, we computed the NDR Score (Methods) and explored its association with TPM levels using data from the GTEX Whole Blood RNA-Seq dataset (Supplementary Figure 1a). We identified a moderate inverse correlation between the NDR Score and gene expression levels (Spearman’s ρ = −0.614, p < 0.05). Notably, we observed distinct clusters of points representing silenced and expressed genes, particularly around NDR Scores of 1.0 and 0.5. A clear separation between expressed and silent genes was evident. However, we observed considerable overlap in mean NDR score across TPM groups of varying expression levels, suggesting limitations in the NDR score’s ability to finely discriminate gene expression levels (Figure 1c).

### GC-bias correction improves NDR signals

GC-bias refers to the correlation between read coverage and GC content of the sequenced region. This bias arises during library preparation, notably in PCR amplification and binding-based purification steps, leading to an underrepresentation of regions with extreme GC content^22^. GC-bias can obscure the desired signals in the analysis of sequencing data related to quantifying of coverage within a genome, and has been applied in cfDNA analysis to improve signals for detection of copy number alterations^23^ and assessment of chromatin accessibility^24^. To account for GC bias in the analysis of NDR scores, we adapted an existing GC-bias estimation method^24^ for the correction of sequencing coverage at the base-pair level. This approach enables the calculation of NDR scores using corrected sequencing coverage (Methods, Figure 1a).

Following GC-bias correction, a robust correlation between original and GC-bias corrected NDR scores (Spearman’s correlation, *R* = 0.91; Figure 1d), indicating that the correction process preserved the underlying biological signal. In addition, consistent trends in relative coverage at NDR sites across varying gene expression levels were maintained, accompanied by a significant reduction in the coverage variability across gene expression groups (Figure 1e, f; Supplementary Figure 1b). Comparing mean NDR scores before and after GC-bias correction, an improved separation of different gene expression groups was observed, attributing to the reduction in variation of NDR scores within each group (Figure 1f). This is reflected by a significant decrease in mean absolute deviation (MAD) of the NDR scores observed post-correction (Figure 1g, *p* < 0.01). Comparison of NDR scores between pre-defined blood- and CRC-specific gene sets (Methods) in CRC and healthy samples shows a slight improvement in the separation of mean NDR scores between the two gene sets in healthy cfDNA samples (Supplementary Figure 1c, d).

### Identification of differential NDR score (DNS)

To detect tissue-specific and treatment-associated changes in NDR scores across samples, we evaluated differences in NDR scores between groups based on sample type and treatment status. Given that NDR score differences follow a normal distribution (Supplementary Figure 2b), a Z-test was implemented for comparative analysis. To improve the accuracy of differential NDR score (DNS) analysis, standard deviation shrinkage was applied to address heteroskedasticity before performing the Z-test (Methods, Figure 2a), a method hereafter referred to as NDRDiff.

**Figure 2.**
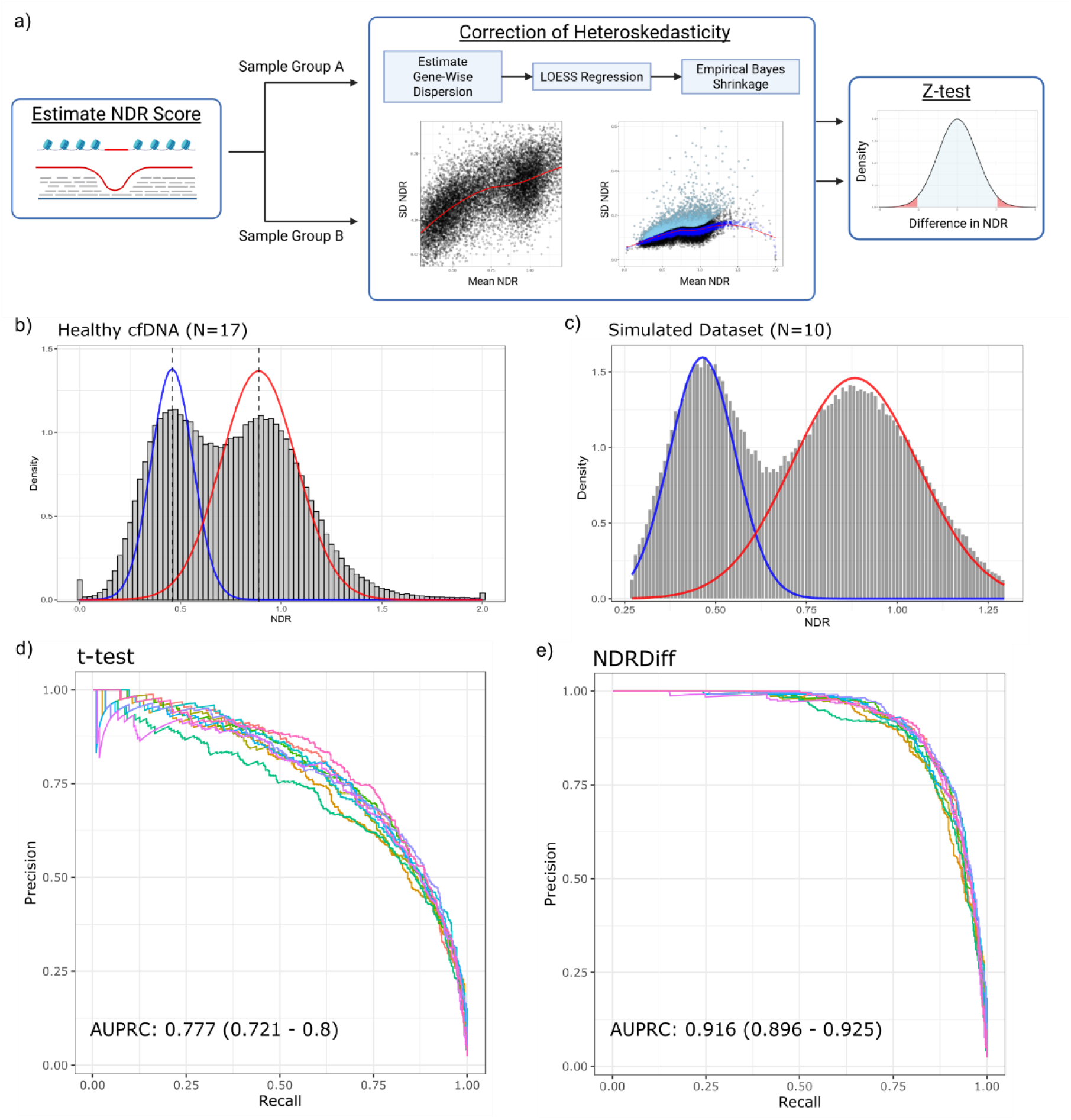
Evaluation of statistical tests for detection of differential NDR score (DNS) genes. **(a)** Schematic of procedure for NDRDiff (Z-test with shrinkage of standard deviation). Created in BioRender. Skanderup, A. (2025) https://BioRender.com/7zlj88e. **(b, c)** Distribution of NDR scores in the (b) healthy cfDNA samples (n=17) and (c) simulated dataset (n=10). **(d, e)** Precision-Recall curve for the detection of significant DNS genes in (d) NDRDiff and (e) t-test.

To evaluate the performance of NDRDiff compared to standard t-test, a simulated dataset was generated for validation. Based on the observation that the overall distribution of NDR scores appear to follow a mixed-effect normal distribution (Figure 2b), we generated 10 simulated datasets using bimodal parameters (Model 1: μ=0.459, σ=0.098, λ=0.354; Model 2: μ=0.886, σ=0.188, λ=0.646; Methods), producing a distribution in the mixed-effect model that mirrors expected profiles (Figure 2c). Each dataset included 500 simulated DNS genes. Comparison of the paired groups in the simulated datasets with NDRDiff identified 532 (range 520-542) DNS genes (FDR < 0.05, |ΔNDR| ≥ 0.2) and demonstrated strong predictive accuracy, with a Area Under Precision-Recall Curve (AUPRC) of 0.916 (range 0.896 – 0.925) (Figure 2e, Table 2). In contrast, the t-test identified 506 (range 486-526) DNS genes (p < 0.01, |ΔNDR| ≥ 0.2), with a lower AUPRC of 0.777 (0.721 – 0.800) (Figure 2d and Table 2). Overall, standard deviation-shrinkage prior to Z-test enhanced the sensitivity for detecting DNS genes relative to t-test while maintaining high specificity.

**Table 2.** Performance metrics of NDRDiff and t-test for differential NDR score (DNS) analysis.

| Performance Metrics | NDRDiff | t-test |  |  |
| --- | --- | --- | --- | --- |
|  | FDR < 0.05 | FDR < 0.05 | p < 0.01 | p < 0.05 |
| **Positive Predictive Value (PPV)** | 0.805 (0.793-0.816) | 0.973 (0.952-0.993) | 0.7 (0.68-0.719) | 0.348 (0.343-0.354) |
| **Negative Predictive Value (NPV)** | 0.996 (0.996-0.997) | 0.977 (0.977-0.978) | 0.993 (0.992-0.993) | 0.999 (0.998-0.999) |
| **Sensitivity** | 0.856 (0.849-0.863) | 0.0776 (0.0573-0.0979) | 0.708 (0.692-0.724) | 0.945 (0.94-0.951) |
| **Specificity** | 0.995 (0.994-0.995) | 1 (1-1) | 0.992 (0.992-0.993) | 0.956 (0.955-0.957) |
| **Significant Genes Detected** | 532 (520-542) | 40 (29-51) | 506 (486-526) | 1357 (1334-1380) |
| **Area Under Precision-Recall Curve (AUPRC)** | 0.916 (0.896 - 0.925) | 0.777 (0.721 - 0.8) |  |  |

### Comparison of Healthy vs CRC samples with NDR score

To assess the feasibility of using NDR scores to identify differential molecular profiles between sample groups, our approach was initially applied to compare healthy (n=17) and mCRC (n=31) samples. A total of 531 DNS genes were identified (FDR< 0.05, |ΔNDR| ≥ 0.2), with 419 genes showing a decrease in NDR scores in mCRC samples (up-regulated in mCRC) and 112 genes showing an increase in NDR scores in mCRC samples (down-regulated in mCRC) (Figure 3a, Supplementary Table 1). The identified DNS genes effectively distinguished healthy donors from mCRC samples, and the degree of separation appeared to be associated with the tumor fraction of the cancer samples (Figure 3b, c). Furthermore, a comparison of the overlapping transcripts from the identified DNS genes with Blood and CRC transcriptomic datasets confirm tissue-specific gene expression of the identified DNS genes (Supplementary Figure 3c).

**Figure 3.**
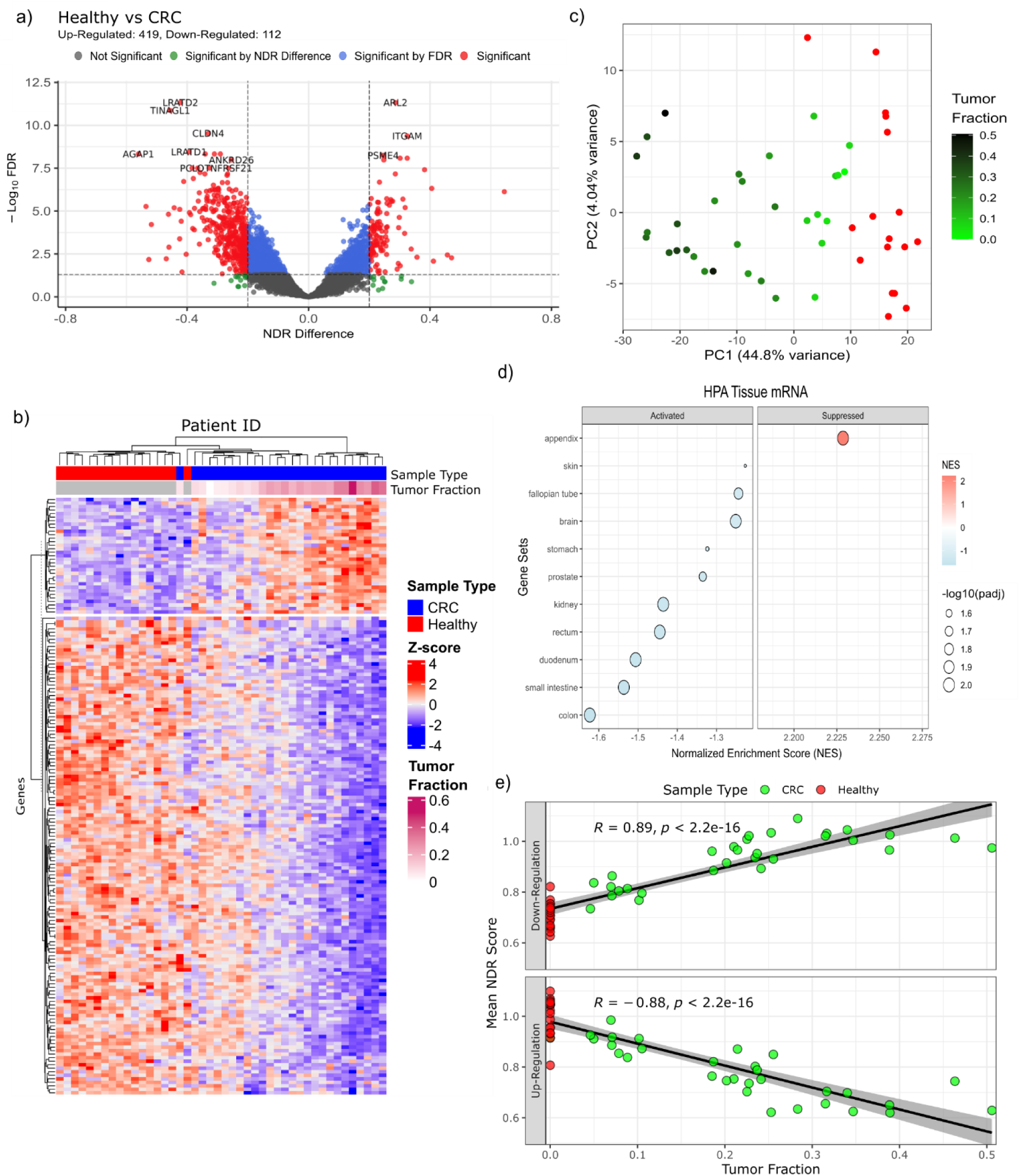
Validation of NDRDiff with comparison of healthy (n=17) and mCRC (n=31) samples. **(a)** Volcano plot of -log10 statistical significance (y-axis) against difference in NDR scores (x-axis) for NDRDiff comparison of healthy and mCRC samples. Significant DNS genes are indicated in red. **(b)** Heatmap of the NDR scores of the identified DNS genes across healthy and mCRC samples. The top annotation indicates the sample type and tumor fraction of the samples. Tumor fraction is not applicable to healthy samples and is greyed out correspondingly. **(c)** PCA plot of healthy (red) and mCRC (green) samples across identified DNS genes. **(d)** GSEA plot depicting the normalized enrichment scores (NES) of top activated and suppressed gene sets identified for HPA tissue mRNA gene sets. Activated gene sets show decrease in NDR scores and suppressed gene sets show increase in NDR scores. **(e)** Scatter plots of mean NDR scores of identified DNS genes against tumor fraction in healthy (red) and mCRC (green) samples. Correlation between mean NDR scores and tumor fraction was examined with Pearson correlation. Top panel: DNS genes down-regulated in mCRC samples, bottom panel: DNS genes up-regulated in mCRC samples.

Among the DNS genes identified are CRC-related genes, such as CLDN4, and blood-related genes, such as BIN2 and IRAG2. CLDN4 is known to be overexpressed in various cancers, including CRC, with a study suggesting that targeting CLDN4 can enhance the anti-tumoral effects of chemotherapeutic agents^25^. Consistent with this, CLDN4 was found to be activated in mCRC samples (ΔNDR=-0.33, Supplementary Figure 3b) compared to blood samples. Conversely, an increase in NDR scores was observed in blood-related genes, such as BIN2 and IRAG2 (Supplementary Figure 3b), in mCRC samples (ΔNDR=0.382, 0.301). BIN2 and IRAG2 are both expressed in platelet cells and involved in platelet activation and aggregation^26,27^.

Gene set enrichment analysis (GSEA) on MSigDB hallmark gene sets^28^ demonstrated suppression of interferon alpha and gamma responses, inflammatory responses, and JAK/STAT signaling pathways in mCRC samples, which are critical for immune activity and regulation in the blood (Supplementary Figure 3d). Conversely, we detected activation of E2F targets, which play a key role in cell cycle progression and are frequently associated with cancer proliferation. GSEA of Human Protein Atlas (HPA) Tissue Gene Expression Profiles gene sets^29^ revealed significant activation of gastrointestinal tissue gene sets in mCRC samples, including those from the colon, small intestine, duodenum, and rectum (Figure 3e, Supplementary Figure 3a). In summary, the identification of DNS genes and enriched gene sets related to CRC and blood cells validates that the comparison of NDR scores can distinguish between cancer and normal samples and retrieve tissue-specific expression profiles.

## Discussion

By analyzing cfDNA NDR coverage profiles, gene expression patterns were inferred. To overcome the limitations of existing methods, GC-bias correction and enhanced statistical approaches were implemented to enrich relevant NDR score signals. The approach was validated using simulated datasets and comparisons between healthy and mCRC samples. This work represents a novel application of cfDNA analysis for the comparative analysis of NDR profiles across sample groups.

The study of cfDNA fragmentation profiles and their association with gene expression has been previously investigated^18,19^. However, earlier analyses were largely confined to inference of gene expression states (expressed or silenced) and estimation of cell type contribution to cfDNA. In this study, we demonstrate that incorporating GC-bias correction and statistical methods accounting for the heteroskedasticity of NDR scores enhances the detection of differential NDR score genes. The genes identified using this approach are shown to be relevant across different contexts, including simulated datasets (Figure 2) and comparison between healthy and cancer samples (Figure 3).

Prior studies on cfDNA analysis have employed gene signatures derived from orthogonal approaches, such as DNA methylation sequencing or RNA sequencing, to evaluate tissue-specific origins of cfDNA^21,30^ and to estimate tumor burden^20^. These studies have demonstrated the capability to specifically identify tissue-specific and tumor-specific cfDNA fragments, marking a notable advancement in the clinical utility of cfDNA. Consistent with these findings, our analysis comparing healthy and cancer samples identified DNS genes that distinguish between the two groups and shows a correlation with tumor fraction in cancer samples (Figure 3). Our approach enables the identification of gene signatures specific to cfDNA NDR profiles, which may improve tracking of tumor contributions across different cell types and cancer types.

The use of cfDNA for inferring gene expression remains in its early stages, and several considerations can be addressed to enhance the accuracy of NDR signals. First, NDR scores were estimated based on relative coverage within a fixed region spanning −50 bp to +150 bp relative to the TSS. However, the actual region of interest, characterized by a dip in the cfDNA profile, may not consistently align with this fixed region. This potential misalignment highlights the need for methods to identify and evaluate optimal NDR sites for each transcript based on their correlation with gene expression. Second, the predominance of cfDNA from blood-derived sources poses a significant challenge, as cancer-specific NDR signals can be obscured by contributions from blood signals. Developing robust approaches to quantify and account for the influence of blood-derived cfDNA will be essential for improving the detection of cancer-specific signals. Lastly, cfDNA analysis primarily requires sequencing of a ±2 kb region around the TSS for each gene, making it feasible to integrate this approach into existing targeted sequencing panels or whole-exome sequencing platforms. Such integration could enable cost-effective evaluation of inferred gene expression and facilitate the broader adoption of cfDNA-based methods in clinical applications.

## Methodology

### Sample Collection

This study was approved by the institutional review boards of SingHealth (2018/2795, 2019/2401), and all procedures were conducted in accordance with ethical guidelines. Written informed consent was obtained from all participants prior to sample collection. Venous blood samples were collected in EDTA tubes to prevent coagulation. Samples were either stored at −80°C for later analysis or processed within two hours of collection to maintain integrity. Whole blood was centrifuged at 300 g x 10 min and 9730 g x 10 min to isolate the plasma layer, which was subsequently stored at −80°C until further analysis.

### Library Preparation

Plasma DNA was extracted using the QIAamp Circulating Nucleic Acid Kit (QIAGEN, 55114) according to the manufacturer’s protocol. Library preparation was conducted with KAPA HyperPrep Kit (Roche, KK8504), using up to 100 ng of cfDNA each sample. Following end repair and A-tailing, the cfDNA was ligated with custom adapters containing a random 8-mer adjacent to the library index site, synthesized by Integrated DNA Technologies (IDT). Post-ligation cleanup was performed using 0.8X Agencourt AMPure XP Beads, and the adapter-ligated cfDNA was eluted in 20 µL of nuclease-free water. Library amplification was carried out using KAPA HiFi HotStart Ready Mix (Roche, KK2602) and 1X Agencourt AMPure XP Beads (Beckman Coulter, A63882), followed by elution of the amplified library in 20 µL of Elution Buffer (10 mM Tris-HCl, pH 8.0). Quantification of the plasma DNA libraries was performed using the KAPA Universal Library Quantification Kit (Roche, KK4824), and the library quality was assessed with Agilent High Sensitivity DNA Kit (Thermo Fisher Scientific, NC1738319). Sequencing was carried out on an Illumina NovaSeq 6000 System, generating 2 × 150 bp paired-end reads. dWGS cfDNA samples from healthy donors were sequenced to ∼60X coverage, whereas those from mCRC patients were sequenced to ∼120X coverage.

### Nucleosome-Depleted Region (NDR) Score Estimation

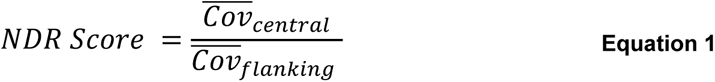

where 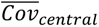 denotes the mean sequencing coverage in the central region surrounding the TSS (−50 to +150 bp relative to the TSS), and 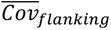 represents the mean coverage in the upstream and downstream flanking regions (−2000 to -1000 bp and +1000 to +2000 bp relative to the TSS).

The NDR score was estimated as described in Equation 1. The mean sequencing coverage of the central region surrounding the transcriptional start site (TSS, -50 to +150 bp relative to the TSS) was normalized by the mean coverage of the flanking regions (−1000 to −2000 bp upstream and +1000 to +2000 bp downstream). Only reads with a mapping quality score of ≥20 was included for the coverage calculation. NDR scores were estimated for selected transcripts in the Matched Annotation from NCBI and EMBL-EBI (MANE) Select set, where transcript at each genomic locus was selected to reflect the biology of the locus^31^.

GC-bias correction was performed prior to NDR score estimation to account for biases arising from variations in GC content across different transcripts. Fragment-length-specific GC-bias estimates were generated using Griffin, with the provided Snakemake workflow file using default parameters (Doebley et al., 2022). Each sequencing read was assigned a GC-bias estimate based on its fragment length and GC content. Bias-corrected coverage was then calculated by applying the reciprocal of the GC-bias factor (1/GC-bias) to ensure normalization of coverage across transcripts. Additionally, NDR sites were excluded if more than half of the samples in either the healthy or colorectal cancer cohorts exhibited an average flank coverage of zero or extreme NDR scores (< 0.1 or > 2).

### Differential Comparison of NDR Scores

To compare nucleosome-depleted region (NDR) scores across groups, such as healthy versus colorectal cancer (CRC) samples, a robust statistical approach was developed to account for observed heteroskedasticity. Specifically, we observed that higher NDR scores were associated with increased standard deviations (SD) (Supplementary Figure 2a), violating the assumption of constant dispersion required for standard t-tests. This violation adversely impacted performance, leading to lower positive predictive value (PPV) and reduced sensitivity across different significance thresholds (Table 2). Additionally, the differences in NDR scores across comparison groups generally followed a normal distribution (Supplementary Figure 2b). Drawing inspiration from RNA-Seq methodologies, such as DESeq2^32^, a shrinkage approach for variance estimation was implemented to address these challenges.

The proposed method, termed NDRDiff, incorporates a Z-test with empirical Bayes shrinkage of the SD to adjust for heteroskedasticity. Initially, SD values were estimated for each gene within the respective sample groups. Observations revealed that the log-transformed SD (logSD) of NDR scores approximated a normal distribution (Supplementary Figure 2c). LOESS regression was performed to model the relationship between the logSD and the mean NDR score. Subsequently, empirical Bayes shrinkage was applied to the logSD values using mean and variance from LOESS regression as prior mean and variance respectively, yielding posterior estimates of the logSD:

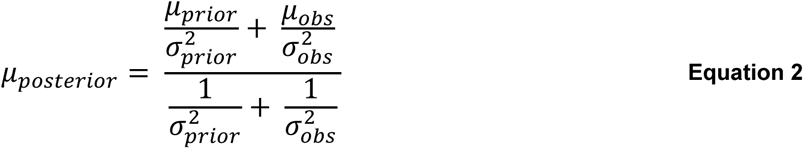

where:

- 𝜇_𝑝𝑜𝑠𝑡𝑒𝑟𝑖𝑜𝑟_ is the posterior (shrunk) estimate of the logSD.
- 𝜇_𝑝𝑟𝑖𝑜𝑟_ and 𝜇_𝑜𝑏𝑠_ are the prior and observed means of logSD, respectively.
- 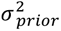 and 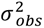 are the corresponding variances.

The shrunk SDs were derived from the posterior estimates of logSD. For genes in which the shrunk SD deviated by more than two SDs above the LOESS regression fit, the original unshrunk SD was retained to avoid false positives due to over-shrinkage. These shrunk SDs were subsequently used in a two-sample Z-test to identify genes with significant differences in NDR scores between sample groups.

### Tissue-Specific Gene Set Selection

Tissue-specific gene sets were identified by comparing gene expression profiles from GTEx Whole Blood and TCGA Colorectal RNA-Seq datasets. Genes were classified as blood-specific if their TPM values were ≥5 in blood and <0.2 in CRCs, and as CRC-specific if their TPM values were ≥5 in CRC and <0.01 in blood. Both datasets were sourced from the UCSC Toil RNA-Seq Recompute Compendium^33^ and accessed via the Xena Functional Genomics Explorer platform (https://xenabrowser.net). The datasets were processed using a standardized workflow to eliminate computational batch effects, ensuring consistency in the analysis.

### Gene Set Enrichment Analysis

Gene Set Enrichment Analysis (GSEA) was performed to explore enriched pathways. GSEA was performed using the fgsea^34^ package with Z-statistics as input, analyzing gene sets from MSigDB Hallmarks^28^ and HPA Tissue mRNA datasets^29^.

### Statistics

Statistical analyses were conducted as follows: group comparisons were performed using the non-parametric Wilcoxon test, which does not assume an underlying data distribution. Correlation analyses were carried out using Pearson correlation when a linear relationship was expected; otherwise, Spearman correlation was applied. For differential NDR score comparisons across sample groups, standard t-tests and Z-tests incorporating dispersion shrinkage (NDRDiff) were employed.

## Supplementary Materials

**Supplementary Figure 1.**
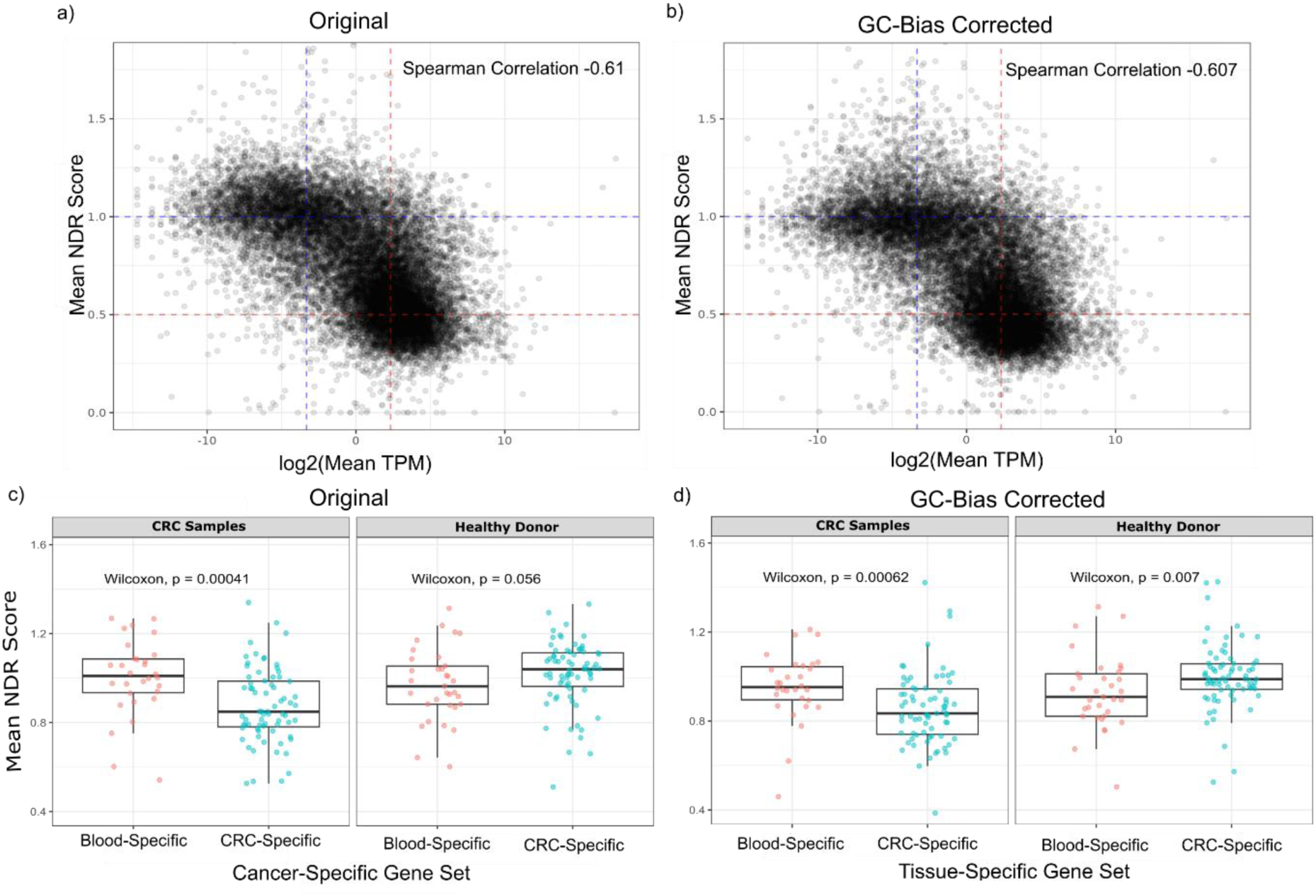
Characterization of GC-bias corrected NDR scores. **(a, b)** Correlation of original (a) and GC-bias corrected (b) NDR scores of healthy cfDNA samples with log2 gene expression from GTEX Whole Blood RNA-Seq. **(c, d)** Comparison of mean NDR score (y-axis) of Blood and CRC-specific gene sets (x-axis) for CRC (left) and healthy samples (right) for original (c) and GC-bias corrected NDR scores (d).

**Supplementary Figure 2.**
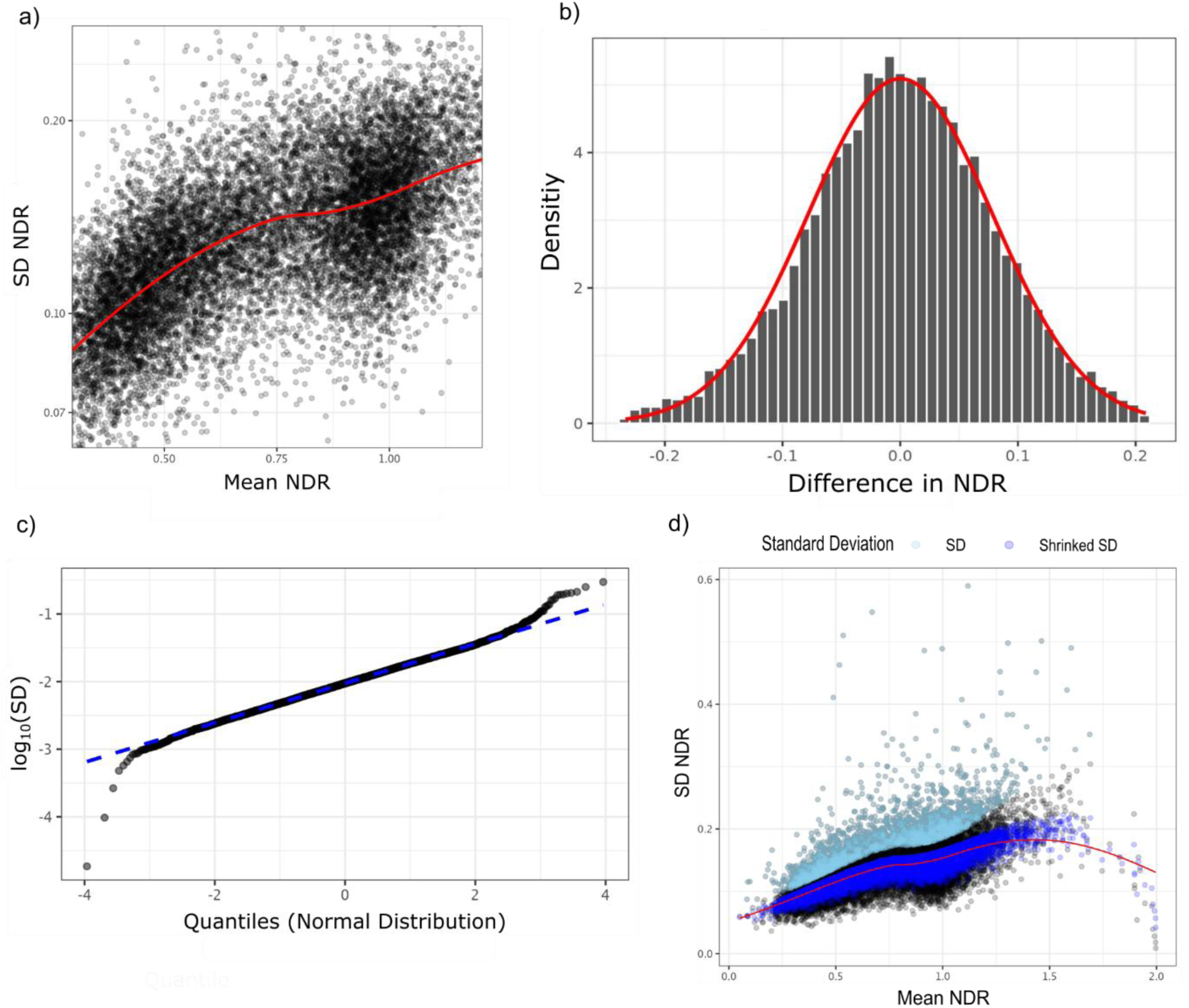
Assessment of NDR score characteristics and distribution for differential comparisons. **(a)** Scatter plots illustrating the distribution of the standard deviation (SD) of NDR scores relative to mean NDR scores. **(b)** Distribution of differences in NDR scores between two sample groups (NDR Diff), with a fitted normal distribution (red line). **(c)** Quantile–quantile plot of log10-transformed SD values. **(d)** Scatter plot showing the relationship between mean NDR scores and SD of NDR scores, with a LOESS regression curve (red line). The blue and light blue dots represent post-shrinkage SD and original SD of transcripts, respectively, with the blue dots indicating the final SD used for the Z-test.

**Supplementary Figure 3.**
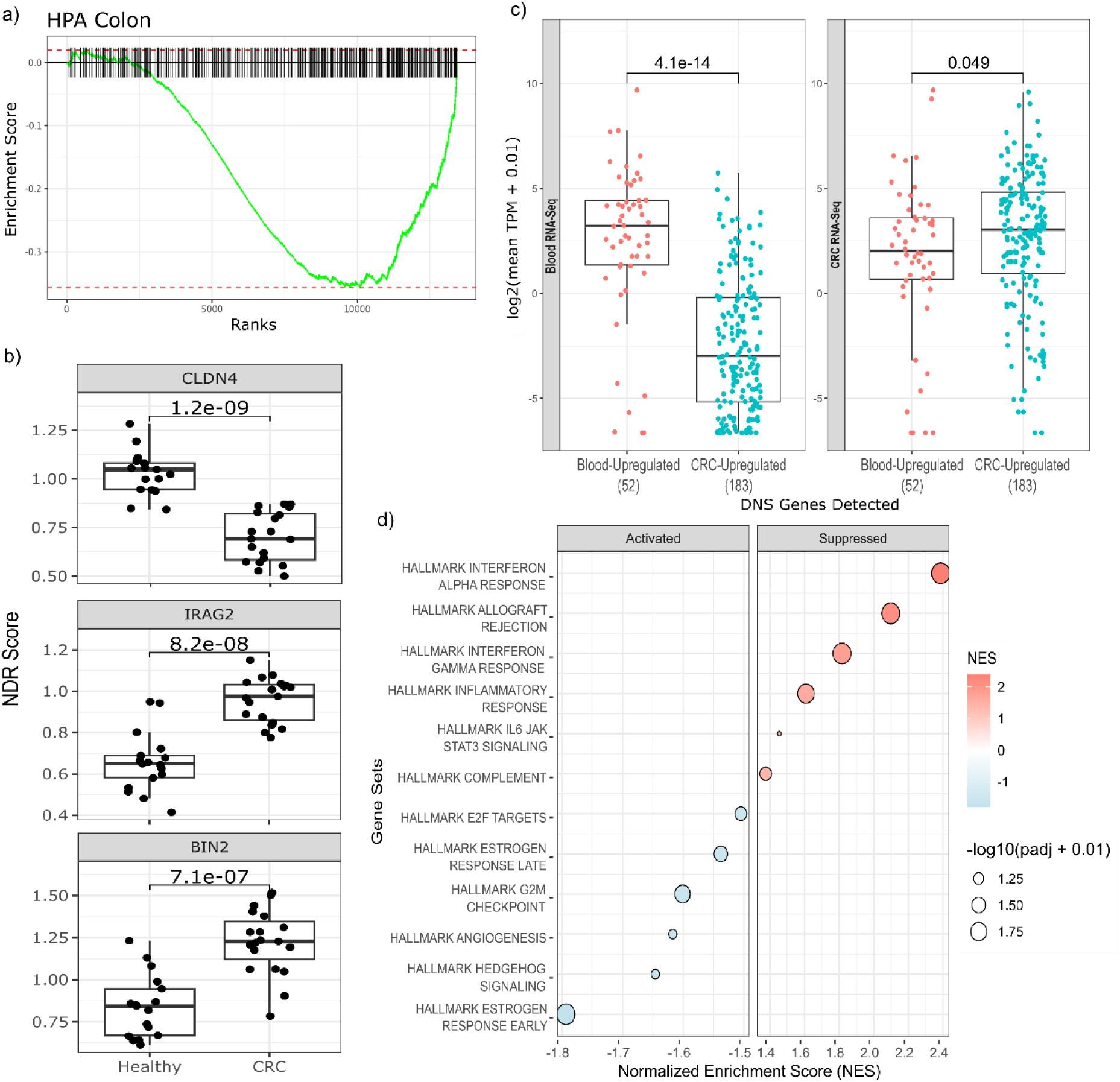
Further validation of NDRDiff results from the comparison of healthy and mCRC samples. **(a)** Enrichment plot for the colon gene set from the HPA Tissue Gene Expression dataset. **(a)** log_2_(TPM + 0.01) values for DNS genes detected in the GTEx Whole Blood RNA-Seq dataset (left) and TCGA COAD-READ RNA-Seq data (right). **(b)** Comparison of mean NDR scores between healthy and CRC samples for selected DNS genes (CLDN4, IRAG2, and BIN2). **(c)** Comparison of mean NDR scores for identified DNS genes that are downregulated (top) and upregulated (bottom) in CRC samples, along with their association with ichorCNA tumor fraction estimates. Healthy samples (assumed to have a tumor fraction of zero) are indicated in red, while cancer samples are shown in green. Pearson correlation was used to assess correlation between mean NDR score and tumor fraction estimates. **(d)** GSEA results showing the normalized enrichment scores (NES) for the significant gene sets from the MSigDB Hallmark database.

## Acknowledgment

Public RNA-Seq from The Cancer Genome Atlas (TCGA) and Genotype-Tissue Expression (GTEx) were accessed from the UCSC Toil RNA-seq Recompute Compendium^33^. TCGA RNA-Seq data used were generated by the TCGA Research Network: https://www.cancer.gov/tcga. The GTEx Project was supported by the Common Fund of the Office of the Director of the National Institutes of Health, and by NCI, NHGRI, NHLBI, NIDA, NIMH, and NINDS.

